# Piggy: A Rapid, Large-Scale Pan-Genome Analysis Tool for Intergenic Regions in Bacteria

**DOI:** 10.1101/179515

**Authors:** Harry A. Thorpe, Sion C. Bayliss, Samuel K. Sheppard, Edward J. Feil

## Abstract

Despite overwhelming evidence that variation in intergenic regions (IGRs) in bacteria impacts on phenotypes, most current approaches for analysing pan-genomes focus exclusively on protein-coding sequences. To address this we present Piggy, a novel pipeline that emulates Roary except that it is based only on IGRs. We demonstrate the use of Piggy for pan-genome analyses of *Staphylococcus aureus* and *Escherichia coli* using large genome datasets. For *S. aureus,* we show that highly divergent (“switched”) IGRs are associated with differences in gene expression, and we establish a multi-locus reference database of IGR alleles (igMLST; implemented in BIGSdb). Piggy is available at https://github.com/harry-thorpe/piggy.

## Background

Whole-genome sequencing has revealed that, in many bacteria, individual strains frequently recruit new genes from a seemingly endless genetic reservoir. The total complement of genes observed across all strains, known as the pan-genome, often numbers tens of thousands, up to an order of magnitude more than the number of genes present in any single genome. In contrast, the “core-genome”, which refers to the complement of genes present in all (or the vast majority) of sampled isolates, can be significantly smaller than the total number of genes in any given genome [1,2]. For example, a study of 328 *Klebsiella pneumoniae* isolates, each of which harbour 4-5,000 genes, revealed a pan-genome of 29,886 genes; only 1,888 (6.8%) of which were universally present (core) [3]. Similarly, genome data for 228 *Escherichia coli* ST131 isolates revealed a pan-genome of 11,401 genes, of which 2,722 (23.9%) were core [4]. The degree of gene content variation in the latter study is particularly striking as these isolates were all from the same sequence type (ST), thus show limited nucleotide divergence in core genes, and are descended from a recent common ancestor.

There is growing recognition that the acquisition of new genes through horizontal gene transfer (HGT) has a central role in ecological adaptation [5]. The emergence and spread of antibiotic resistance, underpinned by the transfer of plasmids and other MGEs, is a pertinent example. The increasing availability of datasets containing thousands of isolates thus offers an unprecedented opportunity for describing the genetic basis of bacterial adaptation. However, the scale of these data presents serious logistic and conceptual challenges in terms of data management and analysis.

Pioneering pan-genome analysis tools, such as PanOCT and PGAP relied on all-vs-all BLAST comparisons between protein sequences, and scaled approximately quadratically with the number of isolates [6,7]. LS-BSR introduced a pre-clustering step which substantially reduced the number of BLAST comparisons, but sacrificed specificity [8]. More recently, the Roary pipeline has rapidly gained in popularity for scalable, user-friendly, pan-genome characterisation [2]. Roary uses a pre-clustering step based on CD-HIT [9], and is more accurate and faster than LS-BSR, meaning that it can analyse 1000s of isolates quickly using modest computing resources.

The concept of the pan-genome, as described above, places an exclusive emphasis on genes; or, more specifically, open reading frames with the potential to encode proteins. This gene-centric perspective has both shaped, and been shaped by, the bioinformatics tools developed to interrogate the pan-genome. For example, Roary works by taking individual protein-coding sequences, pre-defined using Prokka annotation [10], and assigning each to a single cluster of homologous sequences. This approach thus excludes non protein-coding intergenic regions (IGRs) which typically account for approximately 15% of the genome. This is clearly problematic for downstream attempts to identify genotype-phenotype links, as IGRs contain many important regulatory elements including, but not limited to, promoters, terminators, non-coding RNAs, and regulatory binding sites. Moreover, we have recently shown that IGRs are subject to purifying selection in the core-genomes of diverse bacterial species, even when known major regulatory elements are excluded [11,12].

Given that variation in IGRs can have profound phenotypic consequences, it is timely to consider how best to incorporate these sequences into pan-genome analyses. A key question is the degree to which protein-coding genes, and their cognate regulatory elements, should be considered a single “unit”, both selectively (in terms of co-adaptation) and in terms of physical linkage on the chromosome. If physical linkage is assumed to be highly robust, such that genes are mostly transferred along with their cognate IGRs, then in principle the definition of a “gene” could be expanded to include the upstream regulatory regions. On the other hand, if there is moderate or weak linkage between genes and IGRs, such that IGRs can occasionally transfer independently, then the purview of the pan-genome could be expanded to include the full complement of IGR alleles in addition protein-coding sequences.

Consistent with the second model, which allows for independent transfer of IGRs, a landmark study demonstrated that *E. coli* genes can apparently be regulated by alternative IGRs that frequently share no sequence similarity to each other [13]. Moreover, the distribution of these IGRs was incongruent with gene trees, suggesting that recombination can act to replace one IGR with another resulting in regulatory “switches”; a process they call horizontal regulatory transfer (HRT) [13]. It was also noted that conserved flanking genes may facilitate this process by providing localised regions of homology. IGR switches can be accompanied by differential gene expression [13], and may provide a mechanism to offset the fitness costs of harbouring plasmids and other MGEs [4], pointing to a central role for this process in adaptation.

Our current understanding of the evolutionary dynamics of IGRs in the context of bacterial pan-genome leave many open questions. Specifically, it is unclear how IGRs are distributed among isolates within bacterial populations, how commonly IGRs and their cognate genes are co-transferred, or how the frequency of HRT relates to different functional gene categories. A more complete understanding of bacterial adaptation clearly requires a careful consideration of gene presence/absence alongside gene regulation. Here we address this by introducing a new pipeline called Piggy which closely emulates and complements the established pan-genome analysis pipeline Roary [2]. Input and output files for Piggy and Roary use the same format, and run in a similar time on modest computing resources. Piggy provides a means to rapidly identify IGR switches, and more broadly the means to examine the role of horizontal transfer in shaping the bacterial regulome. We demonstrate the utility of Piggy using large genome datasets for single lineages within two bacterial species, both of which are of high public health importance; *Staphylococcus aureus* ST22 (EMRSA-15) and *Escherichia coli* ST131. Conventional pan-genome analyses are applied to analyse and compare core and accessory IGRs / genes in these lineages. In *S. aureus* we show a link between IGR switching and changes in gene expression, and demonstrate proof-of-principle by establishing a multilocus IGR scheme, (igMLST) in BIGSdb [14]. Piggy is available at (https://github.com/harry-thorpe/piggy) under the GPLv3 licence.

## Results

### Staphylococcus aureus ST22

*S. aureus* ST22 (EMRSA-15) is a clinically important hospital-acquired methicillin resistant strain which is common in the UK and is rapidly expanding elsewhere in Europe and globally. Previous work has shown that *S. aureus* ST22 is clonal and has a relatively small set of accessory genes [15,16]. The size of the gene and IGR pan and core-genomes were compared by running 500 ST22 [16] isolate genomes through Roary and Piggy. Frequency histograms and accumulation curves were plotted for both genes and IGRs (Fig 2). The gene-IGR frequency histogram (Fig 2a) shows that there are 2,312 core genes and 1,486 core IGRs, where core is defined as gene presence in > 99% of isolates. The fact that there are fewer core IGRs than core genes is in part due to the exclusion of intra-operonic IGRs < 30 bp. Both distributions conform to the U-shape typically found in such analyses, where the majority of genes/IGRs are either very common or very rare. The gene accumulation curve (Fig 2b) shows a total of 3,225 genes, with a mean of 2,524 genes per isolate. The gradient of the curve is shallow, consistent with the small, closed, pan-genome of clonal ST22 isolates. The IGR curve shows that each isolate has fewer IGRs than genes (1,696 on average per isolate) due to the exclusion of IGRs < 30 bp, but that the total number of IGRs (3,593) is higher than the total number of genes reflecting greater diversity in IGRs than genes. The IGR curve increases more steeply than the gene curve, and does not appear to plateau. Despite these differences, within any given isolate on average 92% of genes and 88% of IGRs were core.

**Fig 1:**
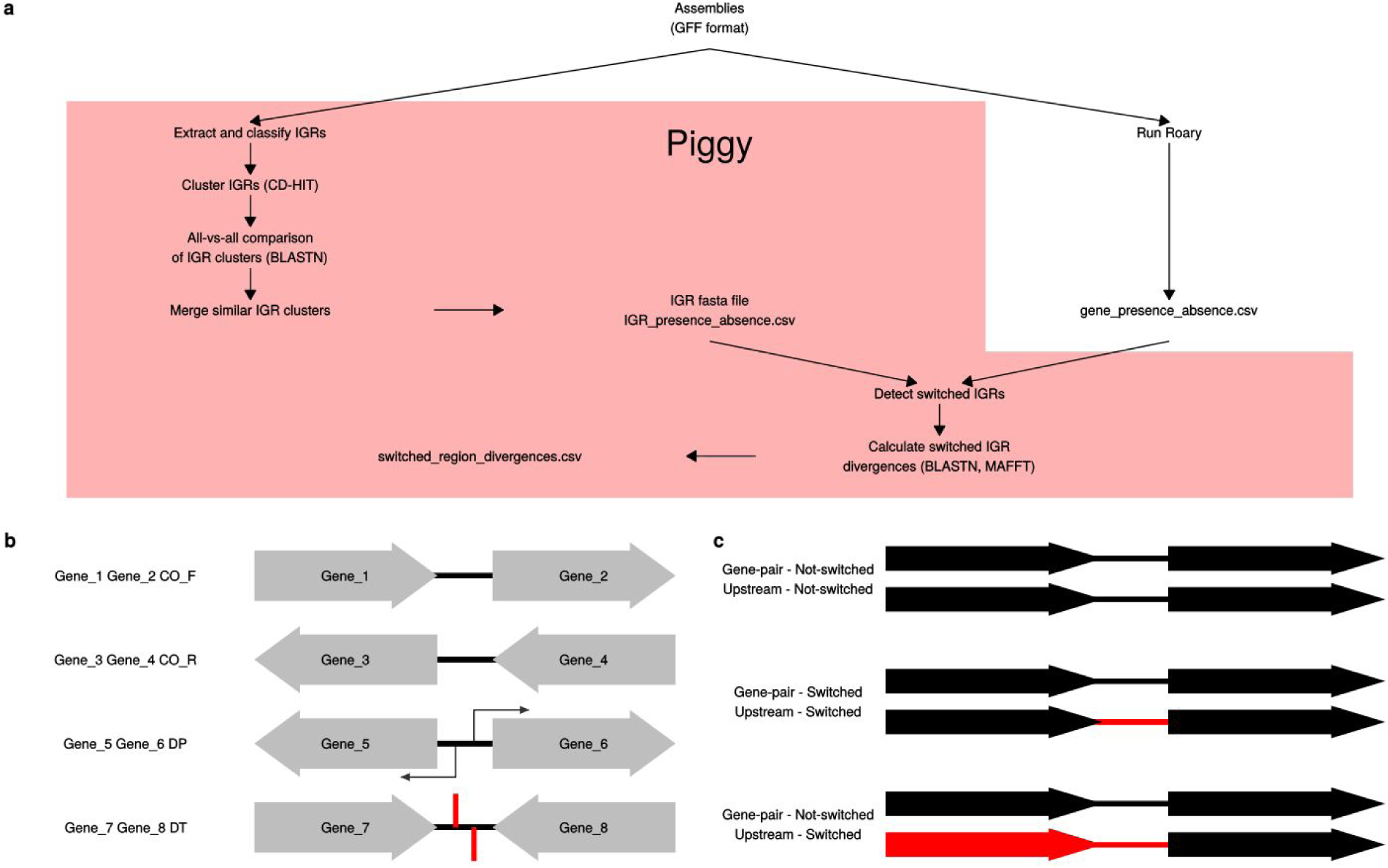
An overview of the Piggy pipeline. a) A schematic to illustrate the Piggy pipeline and how it works alongside Roary. b) IGRs are named according to their flanking genes and their orientations. This naming scheme enables Piggy to link genes with their associated IGRs, and provides information on their orientations. c) A schematic to illustrate the difference between the “gene-pair” and “upstream” methods used to identify candidate switched IGRs.

**Fig 2:**
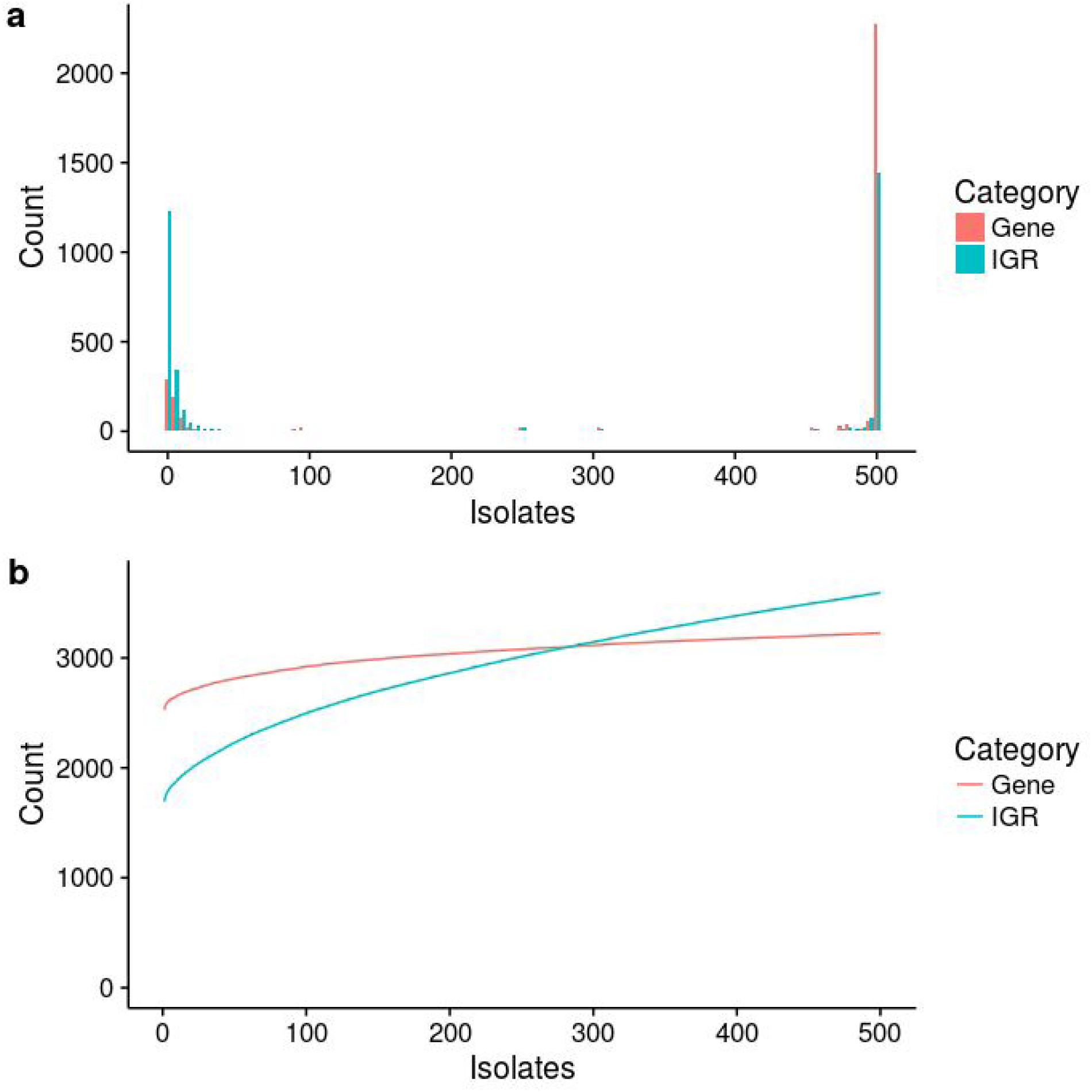
Properties of the *S. aureus* ST22 pan-genome. Genes (red) and IGRs (blue) were analysed. a) Gene and IGR frequency histogram – that is, the number of genes / IGRs present in any given number of isolates. The vast majority of genes / IGRs are either very rare or very common. b) Gene and IGR accumulation curves – that is, the cumulative number of genes / IGRs detected in a given number of isolates.

### Escherichia coli ST131

The utility of Piggy was further validated by re-analysing data from a recent study on the widespread and clinically important *E. coli* lineage ST131 [4]. This dataset contains 236 clinical *E. coli* ST131 isolates from human, domesticated animal, and avian hosts. *E. coli* is a more genetically diverse species than *S. aureus*, and unsurprisingly *E. coli* ST131 has a larger pan-genome than *S. aureus* ST22, with 12,806 genes and 16,429 IGRs (Fig 3a). Of these, 3,285 genes and 1,403 IGRs were core (Fig 3b), out of an average of 4,678 genes and 2,999 IGRs per isolate. Thus despite the differences in diversity, for both *S. aureus* and *E. coli* datasets we found a lower number of core IGRs than core genes, but a high number of accessory IGRs compared to accessory genes. This is illustrated by the fact that the IGR and gene accumulation curves intersect in both species. A lower proportion of both genes (70%) and IGRs (47%) are core within each *E. coli* ST131 isolate, compared to *S. aureus* ST22. Similarly, rare accessory genes and IGRs are much more prevalent in *E. coli* ST131 than in *S. aureus* ST22 with 34% of genes and 55% of IGRs found in < 1% of isolates in *E. coli* ST131, compared with 11% of genes and 40% of IGRs in *S. aureus* ST22.

**Fig 3:**
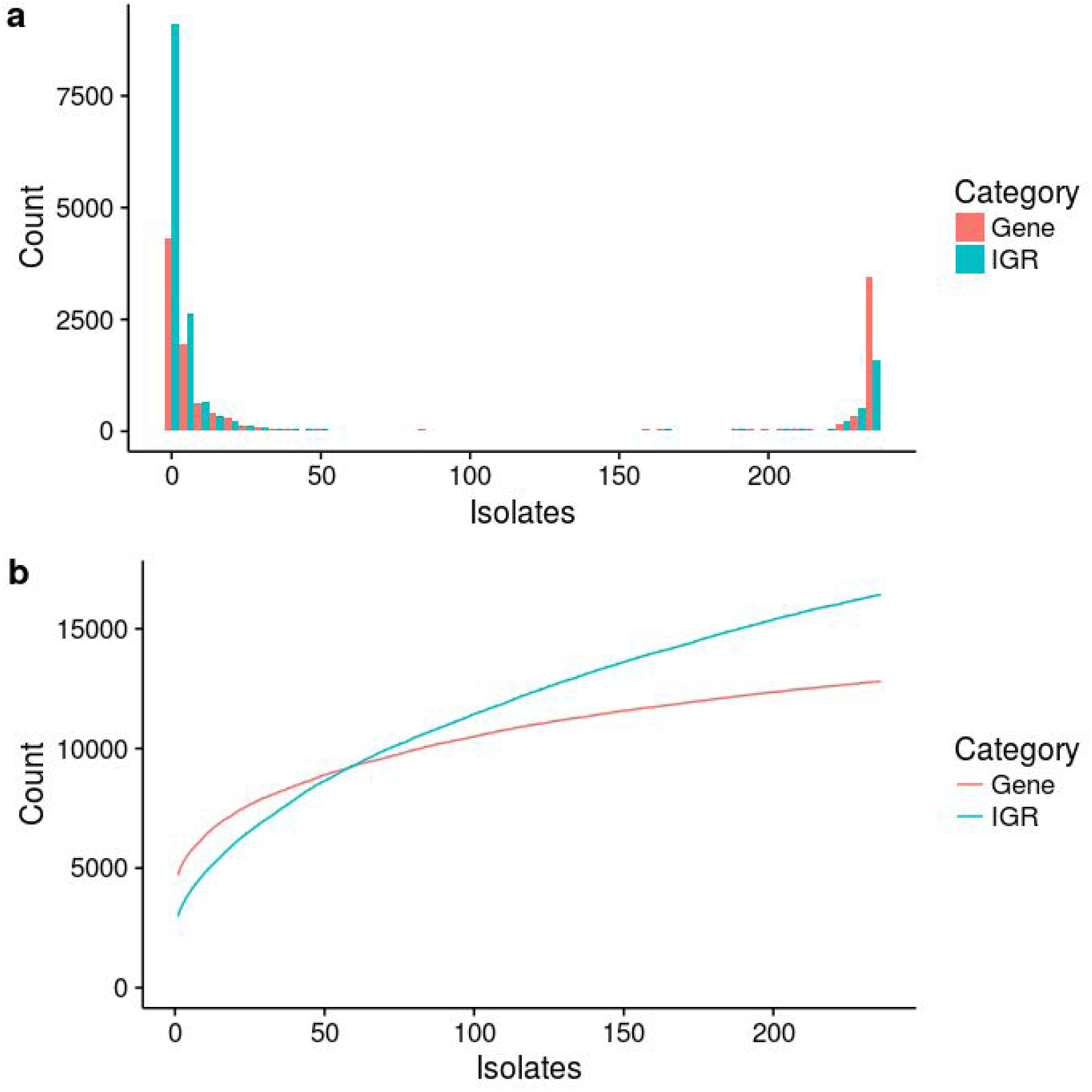
Properties of the *E. coli* ST131 pan-genome. Genes (red) and IGRs (blue) were analysed. a) Gene and IGR frequency histogram – that is, the number of genes / IGRs present in any given number of isolates. The vast majority of genes / IGRs are either very rare or very common. b) Gene and IGR accumulation curves – that is, the cumulative number of genes / IGRs detected in a given number of isolates.

Previous work has found evidence of extensive IGR switching, where the linkage between an IGR and the cognate downstream gene breaks down, resulting in alternative gene / IGR pairs [13]. Piggy provides a list of candidate switching events together for both “gene-pair” and “upstream” approaches (see Methods) at different thresholds of nucleotide identity. For the *E. coli* ST131 data, the pipeline detected 61 cases of putative IGR switching using the most conservative settings (i.e. the conservative gene-pair method, and the alternative IGRs showing no sequence similarity by BLASTN). Relaxing the threshold of sequence identity to < 90% resulted in the identification of an additional 317 candidate switching events, though these possibly reflect either relaxed or positive selection.

### Switched IGRs influence gene expression in *S. aureus*

To examine whether switches in IGRs affect the expression of cognate (downstream) genes, we used a previously published RNA-seq dataset based on four reference *S. aureus* isolates HO_5096_0412 (ST22), Newman (CC8), MRSA252 (CC36), and S0385 (CC398) [17]. Each of these *S. aureus* references isolate represents a distinct major clonal complex, and all were grown under identical conditions with each experiment being replicated. Thus these data provide evidence of the natural variation in gene expression within the *S. aureus* population. By analysing these data alongside the output from Piggy, it is possible to test the extent to which IGR switches between these four genomes can account for the observed variation in gene expression between clonal complexes. First Roary was used to identify a set of 2094 single copy core genes present in all four isolates, and then expression of these core genes was quantified using Kallisto [18]. To do this we used RNA-seq data for two replicates for each of the four reference genomes. The tpm (Transcripts per Kilobase Million) values for each gene are given in Table S1. We then used Sleuth [19] to normalise and filter these counts.

To check the consistency of the data between biological replicates, we first plotted two replicates for each isolate against each other (e.g. Newman replicate 1 vs Newman replicate 2) (Fig 4). These plots were tightly correlated (mean R^2^= 0.98), confirming that the expression values for individual genes were consistent between replicates. We then plotted between-isolate comparisons, again using both replicates for each genome (e.g. Newman replicate 1 vs MRSA252 replicate 1, and Newman replicate 2 vs MRSA252 replicate 2) (Fig 4). These comparisons revealed considerably more scatter, with R^2^ values ranging from 0.76 to 0.9. Given the extremely high R^2^ values for within-isolate comparisons, the decrease in R^2^ for between-isolate comparisons reflects genuine differences in expression between the isolates. We note that a small number of genes show very striking differences in expression between the clonal complexes. For example, the expression of *mepA*, which encodes a multidrug efflux pump, was ∼250 fold higher in Newman compared with the other isolates.

**Fig 4:**
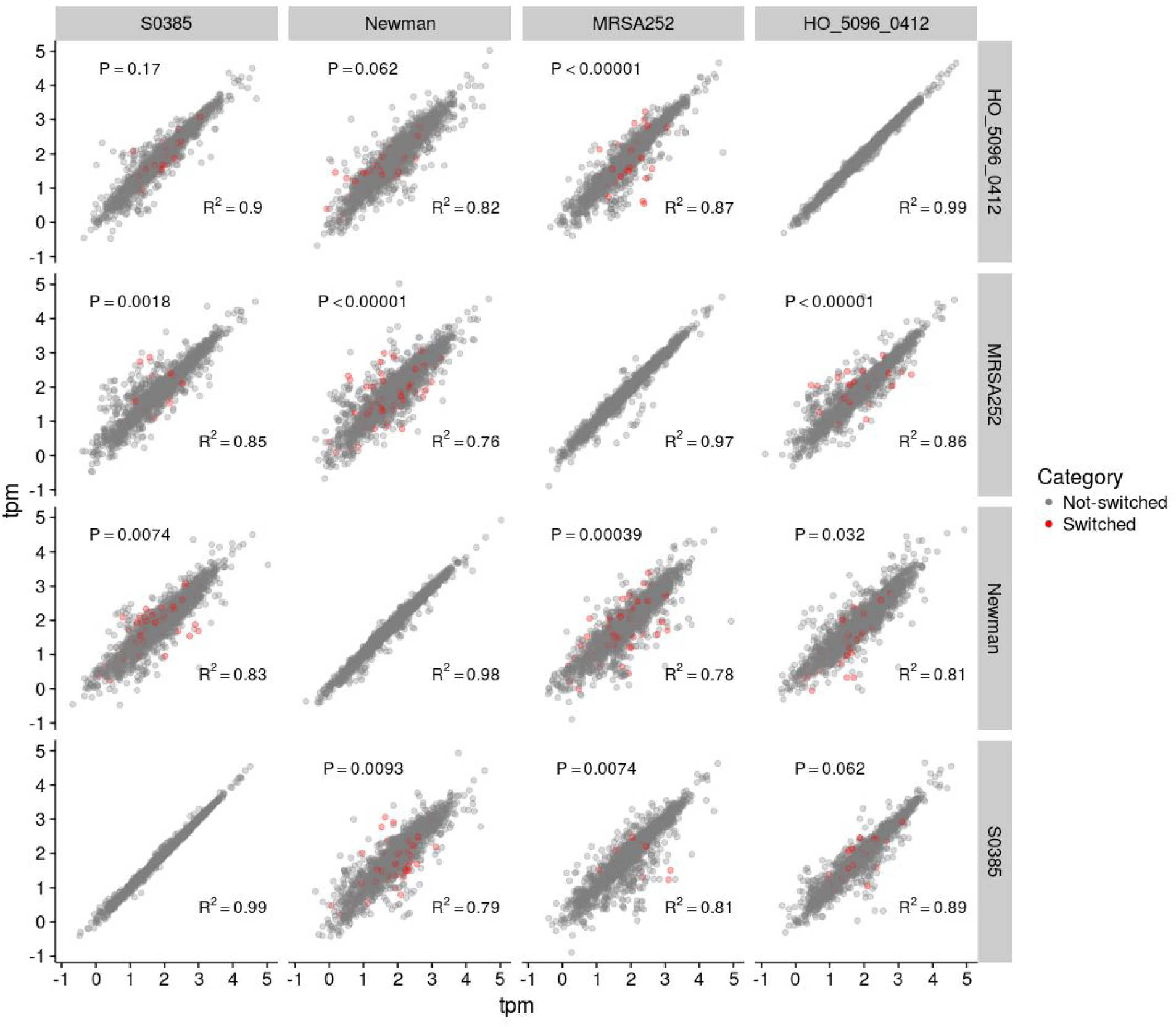
*S. aureus* gene expression data. Pairwise RNA-seq comparisons between four *S. aureus* isolates, where two biological replicates were used for each isolate. The top-left of the diagonal corresponds to comparisons between replicate 1 from different isolates (e.g. SO385 replicate 1 vs HO_5096_0412 replicate 1). The bottom-right of the diagonal corresponds to comparisons between replicate 2 from different isolates (e.g. SO385 replicate 2 vs HO_5096_0412 replicate 2). The diagonal corresponds to comparisons between the two biological replicates from the same isolate. 2094 core genes were analysed in each comparison, and tpm (Transcripts per Kilobase Million) was used to quantify expression. The genes were separated into two categories: Switched (red), and Not-switched (grey), based on their upstream IGRs. The R^2^ value corresponds to all the genes. The P-value corresponds to a Monte Carlo permutation test comparing the residuals of the two groups of genes, where a significant score indicates that the genes downstream of switch IGRs are associated with a higher degree of differential expression (ie higher residuals).

The genomes of each pair of isolates were analysed using Roary and Piggy to identify switched IGRs with a nucleotide identity threshold of < 90% for IGR clusters. For each pair of isolates, we then identified all genes immediately downstream of a switched IGR. As a single switched IGR might impact on the expression of more than one co-transcribed downstream genes we also considered all genes linked in a single operon that could be impacted by a single switching event upstream affecting a shared promoter. For each pair of isolates, we thus identified all core genes putatively affected by upstream IGR switches. We then tested whether these genes showed a higher degree of differential expression by conducting Monte Carlo permutation tests on the residuals from the regressions (Fig 4). For each pairwise comparison of isolates, we summed the residuals of the genes with switched IGRs (shown as red points in Fig 4), and compared this to a distribution obtained by resampling (without replacement) 100,000 random sets of the same number of genes and summing their residuals. We computed a one-tailed p-value by dividing the number of permutations with summed residuals greater than the observed value by 100,000. We then adjusted the p-values using the Benjamini-Hochberg method (Fig 4). Because we used both replicates separately (e.g. Newman replicate 1 vs S0385 replicate 1, and Newman replicate 2 vs S0385 replicate 2), each comparison between pairs of isolates was tested twice independently. In 9/12 pairwise comparisons, the observed residuals of the genes downstream of switched IGRs were significantly greater than expected from the resampled data, indicating that genes with switched IGRs were more differentially expressed than those without. Of the three remaining comparisons, two corresponded to comparisons between HO_5096_0412 and S0385 (P = 0.17, and P = 0.062), and one between HO_5096_0412 and Newman (p = 0.062). The second comparison between HO_5096_0412 and Newman was the most weakly significant result (p = 0.032). Thus, the two replicates for each individual pairwise comparison were largely concordant with each other.

Our analysis confirms that genes downstream of switched IGRs are on average more likely to be differentially expressed than genes not associated with IGR switches as identified using Piggy. To illustrate the genomic context and expression differences of genes with switched IGRs, we selected three of the most differentially expressed genes with IGR switches for the Newman vs MRSA252 comparison, and plotted nucleotide identity across the IGR (calculated as a 20-bp sliding window) alongside gene expression (Fig 5).

**Fig 5:**
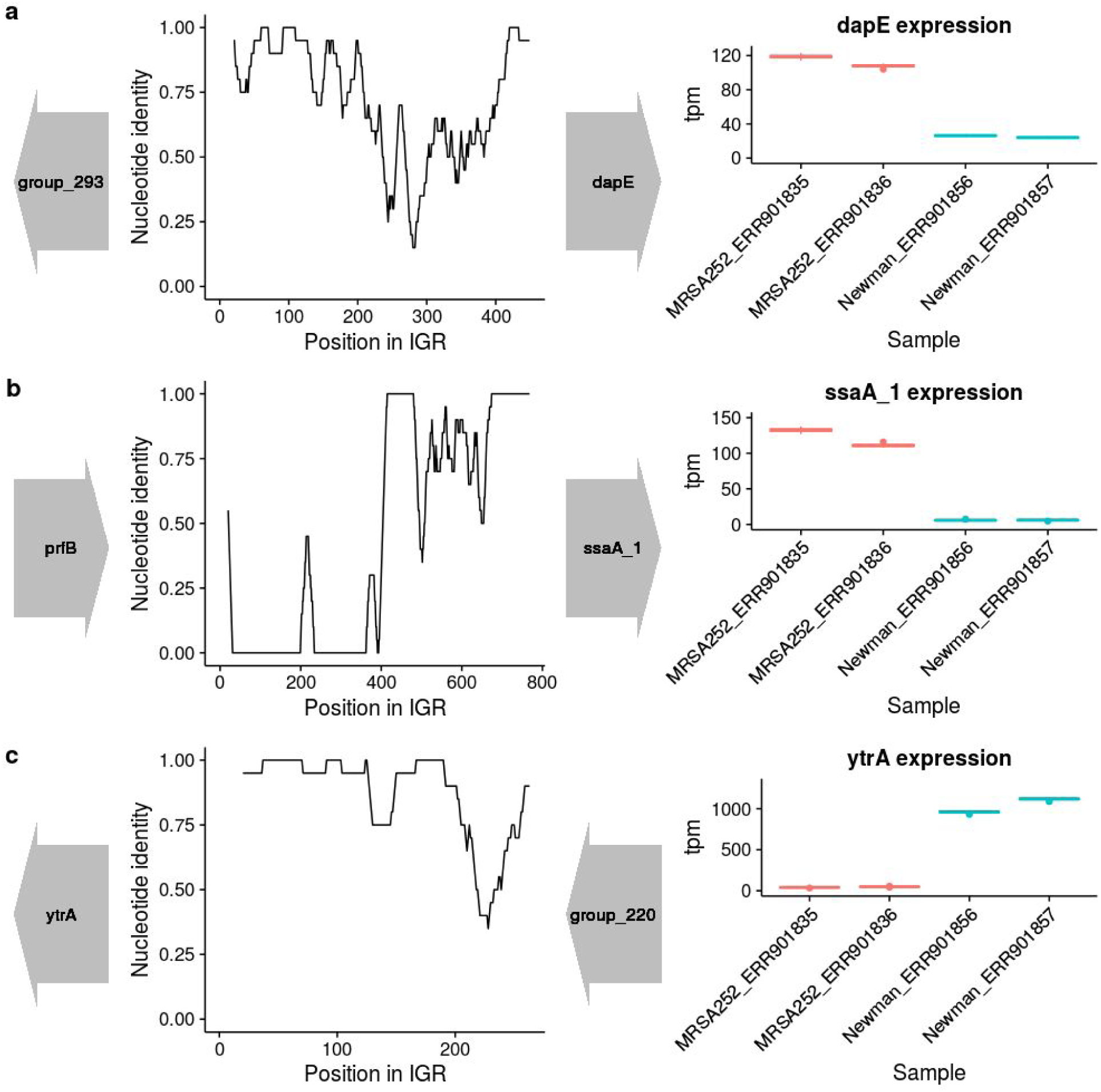
A detailed view of the genomic neighbourhood and expression data for selected genes in Newman vs MRSA252. Nucleotide identity was calculated using a 20 bp sliding window across the IGR, and this is shown alongside the flanking genes in their correct orientation (left). The corresponding expression data for the gene of interest was also shown (right), with the two boxplots per isolate corresponding to the two biological replicates. a) dapE b) ssaA_1 c) ytrA.

### Compatibility and scalability

We have so far demonstrated that Piggy can be used to analyse the intergenic component of the pan-genome and identify IGR switches, and shown that these switches have biological relevance with respect to gene expression. Importantly, Piggy is designed such that the output files are compatible with existing software and databases. The “IGR_presence_absence.csv” file has an identical format to the “gene_presence_absence.csv” file produced by Roary, and can be loaded directly into the interactive browser-based viewer phandango [20] (Fig S1). It can also be used as input, along with a traits file, to Scoary [21] to test for associations between IGRs and phenotypic traits. Moreover, the “representative_clusters_merged.fasta” file can be loaded directly into BIGSdb [14] to create an allele scheme for IGRs. In order to provide proof-of-principle, we created a multilocus IGR (igMLST) scheme in BIGSdb. Briefly, 2631 unique IGR sequences with length ≥ 30bp, from 7 *S. aureus* reference genomes, were entered into the database locus list. Using functionality within the database, these sequences were grouped as a searchable scheme (S_aureus_Intergenic_PIGGY), comparable to MLST, rMLST and wgMLST schemes [22–24]. The distribution of IGRs was analysed for all isolates in the database, identifying IGRs as present in the respective genome if a hit was recorded with nucleotide identity ≥ 70% over ≥ 50% of the sequence using a BLAST word size of 7 bp. The scheme can be found at https://sheppardlab.com/resources/. Finally, Piggy runs in a comparable time to Roary and scales approximately linearly with increasing numbers of isolates, as tested on a MRC-CLIMB [25] virtual machine with 10 vcpus and increasing numbers of *S. aureus* ST22 isolates (Fig S2).

### Discussion

Whole-genome sequence datasets consisting of hundreds or even thousands of bacterial isolates have revealed pan-genomes of many thousands of genes and large differences in gene content between isolates of the same species. Currently, pan-genome diversity is considered almost exclusively in terms of protein-coding genes, despite overwhelming evidence that variation within IGRs impacts on phenotypes. Here we address this by introducing Piggy, a pipeline specifically designed to incorporate IGRs into routine pan-genome analyses by working in close conjunction with Roary [2].

The utility of this approach is demonstrated using large datasets of *S. aureus* ST22 and *E. coli* ST131. Consistent with previous analyses of protein-coding regions [4,15], the IGR component of the ST131 pan-genome (the “panIGRome”) is considerably larger than that for ST22. There was more diversity within IGRs than genes in both species. While some IGRs may be essential for expression of multiple genes, it is expected that IGRs will be subject to less stabilizing selection than protein coding genes [11]. The maintenance of core IGRs in both bacterial genome datasets is consistent with selection acting to conserve them and allows alignment and analysis in much the same way as protein-coding regions.

Variation within regulatory elements located within IGRs can impact on the expression of the downstream gene [13]. Piggy (alongside Roary) provides the means to combine information on genes and their cognate IGRs thus facilitating the detection of “switched” IGRs and downstream genes that are potentially affected. We have shown that in *S. aureus*, genes with switched upstream IGRs show a higher degree of differential expression than those without. This is consistent with previous work on *E. coli* [13], and suggests that the identification of IGR switches using Piggy can provide a useful indication of differential gene expression, even in the absence of RNA-seq data. However, we note that high divergence within IGRs does not necessarily imply selection for differential gene expression, and may instead simply reflect weaker selective constraints. A clear direction for future work is to make constructs consisting of genes with alternative IGRs, in order to directly measure the effect of natural IGR variants on gene expression. Similar experiments have previously been performed in *E. coli* based on variation within promoters [26], and IGRs more broadly [13].

### Conclusions

Driven by recent technical advances in high-throughput sequencing, large whole-genome datasets have provided powerful evidence concerning the genetic determinants that underlie complex multifactorial phenotypes such as virulence. Moreover, associating variation in core and accessory genes with phenotype data is providing new fundamental insight into the ecology and evolution of bacteria. However, in much the same way that non-protein coding DNA in the human genome was initially dismissed as “junk”, omitting IGRs from bacterial genome analysis severely limits our ability to draw inferences on the regulation of gene expression and associated phenotypic consequences. By developing Piggy as an easy-to-use bioinformatics tool with output files that are compatible with existing software and databases (eg Roary, Phandango; Figure S1, Scoary, BIGSdb) we envisage that combined information from genes and their cognate IGRs will vastly improve our understanding of genome evolution in bacteria.

## Methods

### Overview of the Piggy pipeline (Fig 1a)

The first step is to run Roary, as the gene presence absence output file from Roary is used as an input for Piggy. Piggy is then run using the same annotated assemblies as Roary, specifically GFF3 format files such as those produced by Prokka [10]. Piggy extracts intergenic sequences (IGRs) from these files, and uses the flanking gene names and their orientations to name the IGRs (Fig 1b). Including the gene neighbourhood information gives context to the IGR and enables identification of “switched” IGRs. The IGRs are then clustered with CD-HIT [9] at user defined identity thresholds (-n - nucleotide identity, -l - length identity). These two flags allow the user to set the level of stringency for clustering. For example, a conservative approach is to set high values for both nucleotide and length identity such that IGRs must be similar in both nucleotide and length identity to cluster together. By relaxing the length identify whilst maintaining a high nucleotide identity threshold, highly related sequences still cluster even if one is truncated. A representative sequence from each cluster is then used to perform an all-vs-all BLASTN search [27]. This is used to merge similar clusters, which did not cluster with CD-HIT. These clusters are then used to produce an IGR presence absence matrix (“IGR_presence_absence.csv”), in the same format as the gene presence absence matrix (“gene_presence_absence.csv”) produced by Roary. Up until this point, the pipeline is very similar to Roary [2].

### Switched IGR detection

Piggy identifies “switched” IGRs using two methods. The first method identifies adjacent genes on the same contig (gene-pairs), and searches for IGR clusters which lie between these gene-pairs (Fig 1c). Instances where multiple IGR clusters correspond to the same gene-pair are identified as candidate switched IGRs. The second method identifies instances where multiple IGR clusters are upstream of the same gene, which are also putatively switched IGRs. This is a less conservative approach as the downstream gene is not considered in this case, (Fig 1c). The gene-pair method is used by default as it controls against detecting “switching” (recombination) events that encompass more than a single IGR, for example, cases where a mobile element has inserted between two genes. However such cases remain relevant as the regulation of the downstream gene will still be affected.

To ensure that differences in gene annotation between isolates are not erroneously identified as “switching” events, the first and last 30 bp of each flanking gene are searched against the IGRs with BLASTN. Any matches from these searches indicate differences in annotation of gene borders (rather than genuine differences between the IGRs), and these sequences are disregarded. In order to confirm that they represent genuine switching events, candidate switched IGRs are searched against each other with BLASTN with low complexity filtering turned off (-dust no). If there is no significant match they are classed as “switched”, and if there is a significant match they are aligned using MAFFT [28]. The resulting alignment is then used to calculate nucleotide identity (SNPs / shared sites), and length identity (number of shared sites / alignment length). These values can then be used to define an appropriate threshold to identify “switched” IGRs. To aid this, Piggy calculates within-cluster divergences for both genes and IGRs, and these divergences can be used to calibrate Piggy with Roary.

### Datasets

The *S. aureus* ST22 dataset was assembled from published genome sequences of the clinically important lineage ST22 (EMRSA-15) [16] available at http://www.ebi.ac.uk/ena (study number ERP001012). The original genome assemblies were used, and 500 isolates belonging to ST22 were randomly selected for analysis. The *S. aureus* RNA-seq data was previously published [17], and is available at (http://www.ebi.ac.uk/ena, study number ERP009279). This was supplemented with the corresponding reference genomes, HO_5096_0412: HE681097, MRSA252: BX571856, Newman: AP009351, S0385: AM990992, available at (www.ncbi.nlm.nih.gov). The *E. coli* ST131 dataset was also from a previously published study [4], and is available at (http://datadryad.org/resource/doi:10.5061/dryad.d7d71). All complete genomes and assemblies were annotated with Prokka [10].

### Roary and Piggy parameter settings

Roary [2] was run using default parameters except for the following: -e -n (to produce alignments with MAFFT [28]); -i 90 (lower amino acid identity than the default); -s (to keep paralogs together); -z (to keep intermediate files). Piggy was run using default parameters except for --len_id, which controls the percentage of IGR sequences which must share similarity in order to be clustered together. For the *S. aureus* ST22 and *E. coli* ST131 datasets, Piggy was run twice, once with --len_id 10 and once with --len_id 90. The former was used for the pan-genome comparisons between genes and IGRs (Figs 2 and 3) in order to be comparable with Roary (as genes which are clustered together by Roary have a minimum length of 120 bp, and frequently vary greatly in length). The latter was used whenever “switched” IGRs were detected, as this enabled more control over downstream filtering of these sequences.

### RNA-seq analysis

Two biological replicates for each isolate were analysed. Kallisto [18] was used to quantify transcripts (--kmer-size 31 and --bootstrap-samples 100), and Sleuth [19] was used to normalise and filter the counts produced by Kallisto. These counts were then log_10_ transformed, and major axis (MA) regression was performed. Rockhopper2 [29] was used to produce an operon map for each strain by grouping adjacent genes with similar expression profiles together into operons.

### Statistical analysis

All statistical analysis was performed within R version 3.3.2 (https://www.r-project.org). All plotting was performed with ggplot2 [30].

### Declarations

Ethics approval and consent to participate: Not applicable Consent for publication: Not applicable

Availability of data and material: The *S. aureus* ST22 dataset was assembled from published genome sequences of the clinically important lineage ST22 (EMRSA-15) [16] available at http://www.ebi.ac.uk/ena (study number ERP001012). The *S. aureus* RNA-seq data was previously published [17], and is available at (http://www.ebi.ac.uk/ena, study number ERP009279). This was supplemented with the corresponding reference genomes, all available at (www.ncbi.nlm.nih.gov), HO_5096_0412: HE681097, MRSA252: BX571856, Newman: AP009351, S0385: AM990992. The *E. coli* ST131 dataset [4] is available at (http://datadryad.org/resource/doi:10.5061/dryad.d7d71).

Piggy is available at (https://github.com/harry-thorpe/piggy) under the GPLv3 licence.

## Authors’ contributions

HAT designed and implemented the pipeline, and carried out the majority of the analyses, with input from EJF, SCB and SKS. HAT and EJF wrote the manuscript with input from SKS and SCB.

## Acknowledgements

We are very grateful to Torsten Seemann, Andrew Page and João Carriço for encouragement and helpful feedback. We are also grateful to Matt Holden for provision of the *S. aureus* RNA-seq data, to Sandra Reuter for help with the *S. aureus* ST22 data, and to Alan McNally for the *E. coli* ST131 data. This work also greatly benefitted from access to the Medical Research Council funded Cloud Infrastructure for Microbial Bioinformatics (MRC-CLIMB).

---

**Fig S1:**
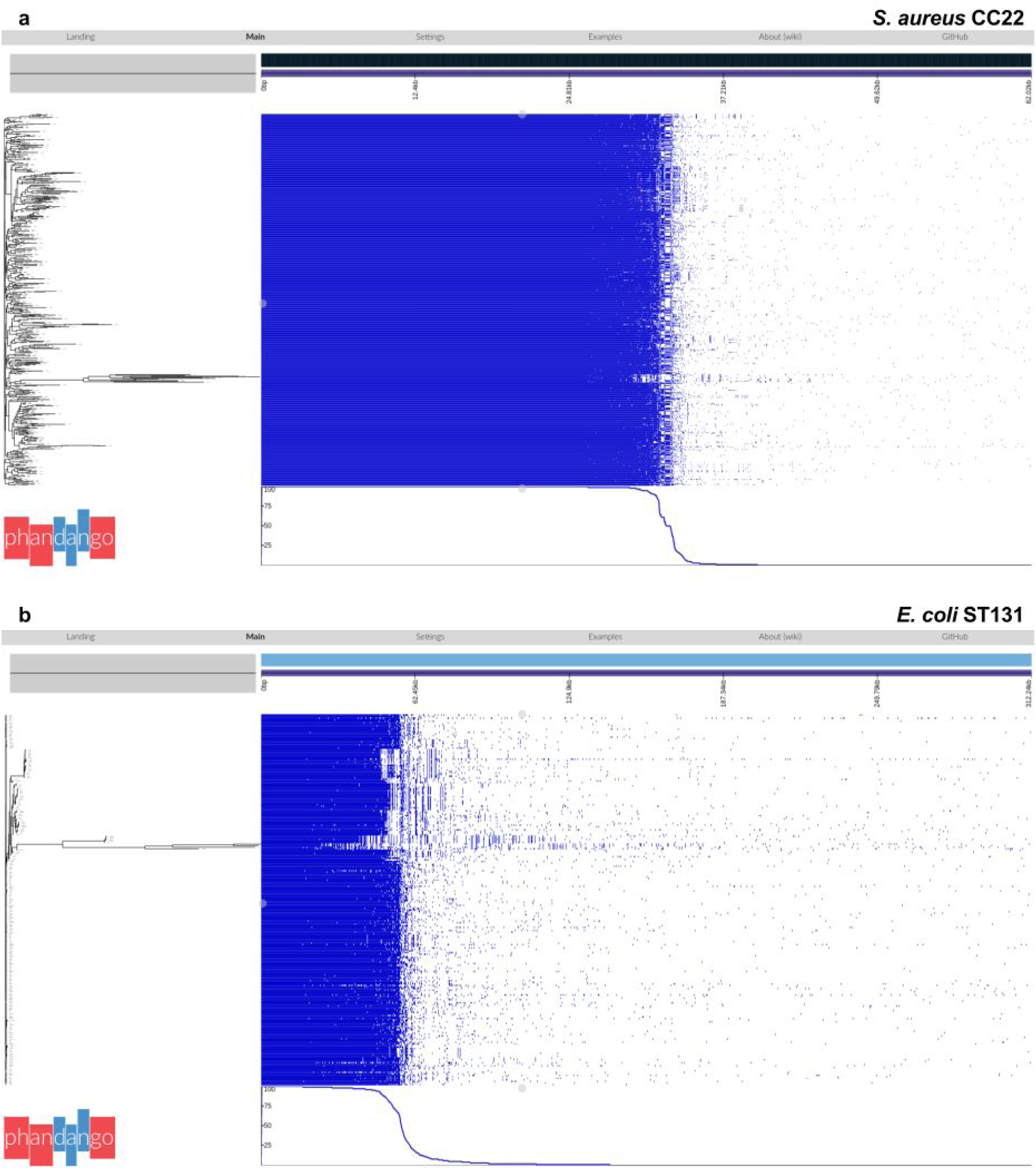
The IGR pan-genome (“panIGRome”) as visualised using Phandango. A neighbour-joining phylogenetic tree was imported into Phandango alongside the IGR_presence_absence.csv file. Each row corresponds to an isolate, and each column corresponds to an IGR, with the IGRs ordered from the left in order of decreasing frequency within the sample. The line graph at the bottom shows the frequency of the IGRs within the sample. a) *S. aureus* ST22 b) *E. coli* ST131.

**Fig S2:**
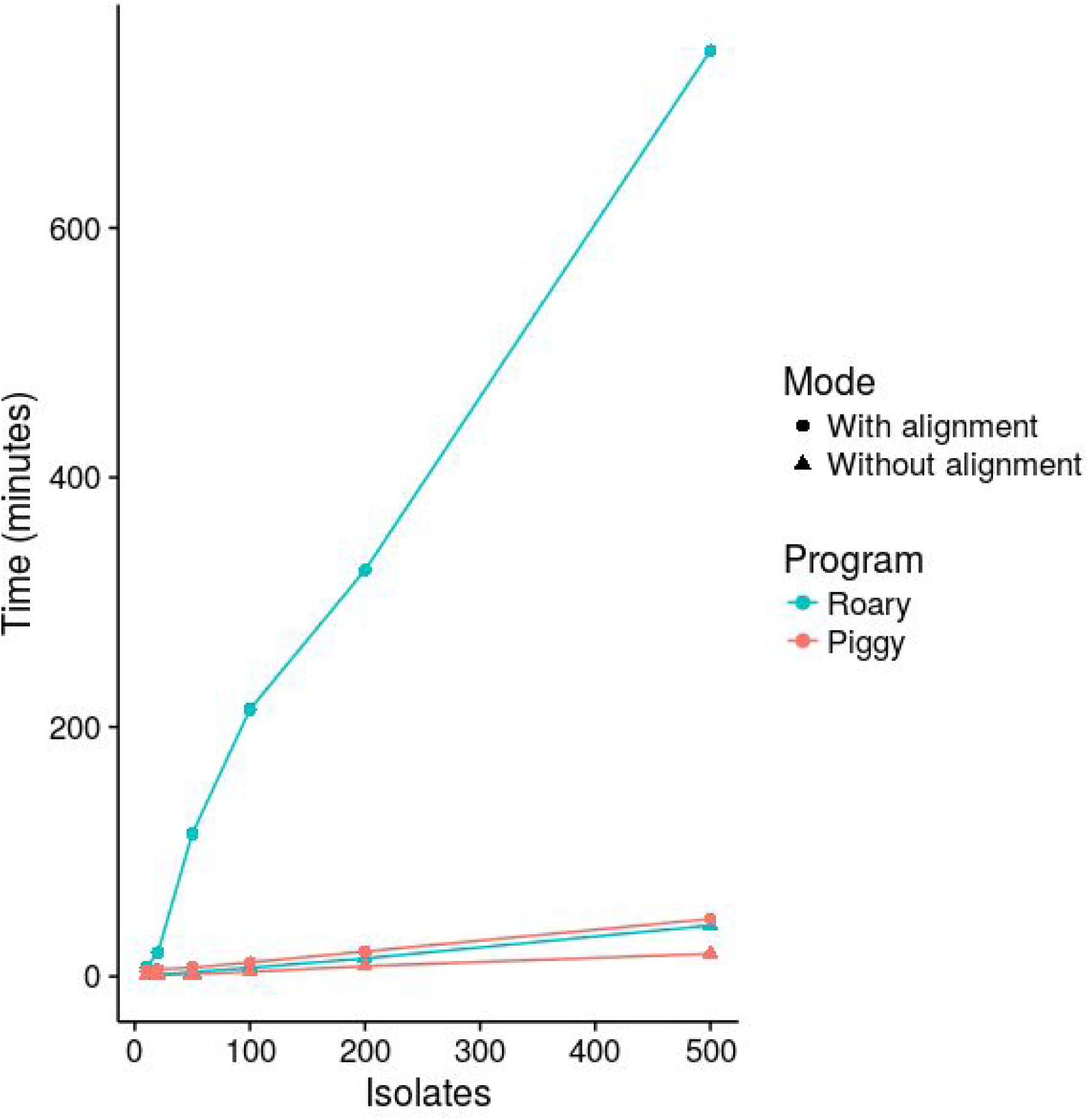
Comparison with Roary. Roary (blue) and Piggy (red) were both run on increasing numbers of *S. aureus* ST22 isolates on a CLIMB virtual machine with 10 vcpus. The programs were both run with (circles) and without (triangles) alignment options.

## References

1. Medini D, Donati C, Tettelin H, Masignani V, Rappuoli R. The microbial pan-genome. Curr. Opin. Genet. Dev. 2005;15:589–94.

2. Page AJ, Cummins CA, Hunt M, Wong VK, Reuter S, Holden MTG, et al. Roary: rapid large-scale prokaryote pan genome analysis. Bioinformatics. 2015;31:3691–3.

3. Holt KE, Wertheim H, Zadoks RN, Baker S, Whitehouse CA, Dance D, et al. Genomic analysis of diversity, population structure, virulence, and antimicrobial resistance in Klebsiella pneumoniae, an urgent threat to public health. Proc. Natl. Acad. Sci. U. S. A. 2015;112:E3574–81.

4. McNally A, Oren Y, Kelly D, Pascoe B, Dunn S, Sreecharan T, et al. Combined Analysis of Variation in Core, Accessory and Regulatory Genome Regions Provides a Super-Resolution View into the Evolution of Bacterial Populations. PLoS Genet. 2016;12:e1006280.

5. Vos M, Hesselman MC, te Beek TA, van Passel MWJ, Eyre-Walker A. Rates of Lateral Gene Transfer in Prokaryotes: High but Why? Trends Microbiol. 2015;23:598–605.

6. Fouts DE, Brinkac L, Beck E, Inman J, Sutton G. PanOCT: automated clustering of orthologs using conserved gene neighborhood for pan-genomic analysis of bacterial strains and closely related species. Nucleic Acids Res. 2012;40:e172.

7. Zhao Y, Wu J, Yang J, Sun S, Xiao J, Yu J. PGAP: pan-genomes analysis pipeline. Bioinformatics. 2012;28:416–8.

8. Sahl JW, Caporaso JG, Rasko DA, Keim P. The large-scale blast score ratio (LS-BSR) pipeline: a method to rapidly compare genetic content between bacterial genomes. PeerJ. 2014;2:e332.

9. Fu L, Niu B, Zhu Z, Wu S, Li W. CD-HIT: accelerated for clustering the next-generation sequencing data. Bioinformatics. 2012;28:3150–2.

10. Seemann T. Prokka: rapid prokaryotic genome annotation. Bioinformatics. 2014;30:2068–9.

11. Thorpe HA, Bayliss SC, Hurst LD, Feil EJ. Comparative Analyses of Selection Operating on Non-translated Intergenic Regions of Diverse Bacterial Species. Genetics [Internet]. 2017; Available from: http://dx.doi.org/10.1534/genetics.116.195784

12. Molina N, Van Nimwegen E. Universal patterns of purifying selection at noncoding positions in bacteria. Genome Res. 2008;18:148–60.

13. Oren Y, Smith MB, Johns NI, Kaplan Zeevi M, Biran D, Ron EZ, et al. Transfer of noncoding DNA drives regulatory rewiring in bacteria. Proc. Natl. Acad. Sci. U. S. A. 2014;111:16112–7.

14. Jolley KA, Maiden MCJ. BIGSdb: Scalable analysis of bacterial genome variation at the population level. BMC Bioinformatics. 2010;11:595.

15. Holden MTG, Hsu L-Y, Kurt K, Weinert LA, Mather AE, Harris SR, et al. A genomic portrait of the emergence, evolution, and global spread of a methicillin-resistant Staphylococcus aureus pandemic. Genome Res. 2013;23:653–64.

16. Reuter S, Török EM, Holden MTG, Reynolds R, Raven KE, Blane B, et al. Building a genomic framework for prospective MRSA surveillance in the United Kingdom and the Republic of Ireland. Genome Res. [Internet]. 2015; Available from: http://genome.cshlp.org/content/early/2015/12/15/gr.196709.115.abstract

17. Warne B, Harkins CP, Harris SR, Vatsiou A, Stanley-Wall N, Parkhill J, et al. The Ess/Type VII secretion system of Staphylococcus aureus shows unexpected genetic diversity. BMC Genomics. 2016;17:222.

18. Bray NL, Pimentel H, Melsted P, Pachter L. Near-optimal probabilistic RNA-seq quantification. Nat. Biotechnol. 2016;34:525–7.

19. Pimentel H, Bray NL, Puente S, Melsted P, Pachter L. Differential analysis of RNA-seq incorporating quantification uncertainty. Nat. Methods. 2017;14:687–90.

20. Hadfield J, Croucher NJ, Goater RJ, Abudahab K, Aanensen DM, Harris SR. Phandango: an interactive viewer for bacterial population genomics [Internet]. bioRxiv. 2017 [cited 2017 Mar 23]. p. 119545. Available from: http://biorxiv.org/content/early/2017/03/22/119545.full.pdf+html

21. Brynildsrud O, Bohlin J, Scheffer L, Eldholm V. Rapid scoring of genes in microbial pan-genome-wide association studies with Scoary. Genome Biol. 2016;17:238.

22. Maiden MCJ, Jansen van Rensburg MJ, Bray JE, Earle SG, Ford SA, Jolley KA, et al. MLST revisited: the gene-by-gene approach to bacterial genomics. Nat. Rev. Microbiol. 2013;11:728–36.

23. Jolley KA, Bliss CM, Bennett JS, Bratcher HB, Brehony C, Colles FM, et al. Ribosomal multilocus sequence typing: universal characterization of bacteria from domain to strain. Microbiology. 2012;158:1005–15.

24. Sheppard SK, Jolley KA, Maiden MCJ. A gene-by-gene approach to bacterial population genomics: Whole genome MLST of Campylobacter. Genes. 2012;3:261–77.

25. Connor TR, Loman NJ, Thompson S, Smith A, Southgate J, Poplawski R, et al. CLIMB (the Cloud Infrastructure for Microbial Bioinformatics): an online resource for the medical microbiology community. Microbial Genomics [Internet]. Microbiology Society; 2016 [cited 2016 Nov 1];2. Available from: http://mgen.microbiologyresearch.org/content/journal/mgen/10.1099/mgen.0.000086

26. Shimada T, Yamazaki Y, Tanaka K, Ishihama A. The whole set of constitutive promoters recognized by RNA polymerase RpoD holoenzyme of Escherichia coli. PLoS One. 2014;9:e90447.

27. Camacho C, Coulouris G, Avagyan V, Ma N, Papadopoulos J, Bealer K, et al. BLAST+: architecture and applications. BMC Bioinformatics. 2009;10:421.

28. Katoh K, Standley DM. MAFFT multiple sequence alignment software version 7: improvements in performance and usability. Mol. Biol. Evol. 2013;30:772–80.

29. Tjaden B. De novo assembly of bacterial transcriptomes from RNA-seq data. Genome Biol. 2015;16:1.

30. Wickham H. Ggplot2: Elegant Graphics for Data Analysis. 2nd ed. Springer Publishing Company, Incorporated; 2009.

